# AlphaFold 3 proteomic modelling reveals multiple photosystem structural homologs in freshwater cyanophages

**DOI:** 10.1101/2024.12.13.628456

**Authors:** Isaac Meza-Padilla, Andrew C. Doxey, Jozef I. Nissimov

**Affiliations:** Department of Biology, University of Waterloo, 200 University Avenue West, Waterloo, Ontario, Canada

## Abstract

Accurate protein structure prediction followed by structural homology detection enable the functional annotation of large numbers of otherwise obscure viral protein-coding genes. Here we employ AlphaFold 3 modeling and DALI structural homology search to predict the structures and functions of all 219 cryptic open reading frames in two representative freshwater cyanophages. We discover 28 previously unknown structural homologs, including three putative PsaD proteins in the same cyanophage, several viral structural proteins, and (to our knowledge) the first reported virus-encoded cyanobacteriochrome (DALI Z-score > 4.2 in all cases). Our results suggest that photosystem proteins may be more widespread in freshwater cyanophages than previously thought and emphasize the importance of applying structural homology detection methods when annotating viral genomes.

## Main

Viruses are probably the most abundant biological systems on Earth^1^. Not surprisingly, the virosphere most likely comprises the largest repertoire of genetic diversity on the planet^2,3^. Coupled to the poor coverage of viruses in genomic databases, such an enormous diversity renders the functional annotation of viral genomes extremely challenging^4^. In many cases, protein-coding genes with no sequence-based homologs in genomic databases can comprise more than 90% of a microbial virus genome^5^. A single viral genome can contain hundreds of protein-coding genes, and thanks to the development of high-throughput metagenomic methods, astronomical numbers of viruses are continuously being discovered^6^. The revolutionary, artificial intelligence-based bioinformatics tool AlphaFold is able to computationally model the three-dimensional structure of proteins with unprecedented accuracy and efficiency^7^. Since the structure of proteins is more conserved compared to their amino acid sequence, structural modelling of viral proteins followed by structural homology search methods makes it possible to predict the function of large numbers of otherwise cryptic proteins at an unprecedented scale^8,9^.

Photosystem protein-coding genes, horizontally acquired from their hosts, are widespread in marine cyanophages and play a fundamental role during infection^10^. Briefly, these proteins help to maintain the photosynthetic efficiency of cyanobacterial virocells^10^. Freshwater cyanophages would be expected to benefit as much as their marine counterparts from maintaining the photosynthetic efficiency of their hosts. In fact, sequence-based homologs of viral photosystem proteins have been detected in environmental samples from freshwater bodies^11^. Curiously, however, a homolog of photosystem genes (*psbA* coding for photosystem II protein D1) has been reported in only one isolated freshwater cyanophage to date^12^.

The aim of this study was to search for cryptic photosystem protein-coding genes in previously isolated freshwater cyanophages. To do this, we conducted proteome structural modelling for the genomes of two representative freshwater cyanophages using AlphaFold 3 (AF3)^7^: CrV-01T infecting the bloom-forming, potentially toxic, nitrogen-fixer *Raphidiopsis raciborskii* (previously *Cylindrospermopsis raciborskii*) Cr2010^13^; and Ma-LMM01 infecting the toxic, bloom-forming, non-nitrogen-fixer *Microcystis aeruginosa* NIES-298^14^. We predicted the structure of all 219 open reading frames (ORFs) with no functionally annotated sequence-based homologs in the NCBI non-redundant protein sequences database. These cryptic protein-coding genes represented 71.5% and 71.2% of CrV-01T and Ma-LMM01 ORFs, respectively. We then used the DALI search tool^15^ to query the PDB and identify structural homologs.

This structural bioinformatics pipeline revealed three structural homologs of the photosystem I reaction center subunit II (PsaD) protein in CrV-01T (ORFs 76, 78 and 115; Fig. 1a,b). Two of them, ORFs 76 and 115, had an identical amino acid sequence. Remarkably, *a posteriori* BLASTP search queried with ORF 76 lead to the sequenced-based identification of another PsaD structural homolog in the CrV-like cyanophage Cr-LKS4 (ORF 79; Fig. 1c), isolated and sequenced by Kolan et al^16^. To the best of our knowledge, this is the first time that putative photosystem I proteins have been reported in isolated freshwater cyanophages. Furthermore, we detected structural homologs for 13 other CrV-01T protein-coding genes, including a Cas-like protein (ORF 58), several viral structural proteins and, to our knowledge, the first reported virus-encoded putative cyanobacteriochrome (CBCR; ORF 65; Fig. 1d-n; Extended Data Table 1).

**Fig 1.**
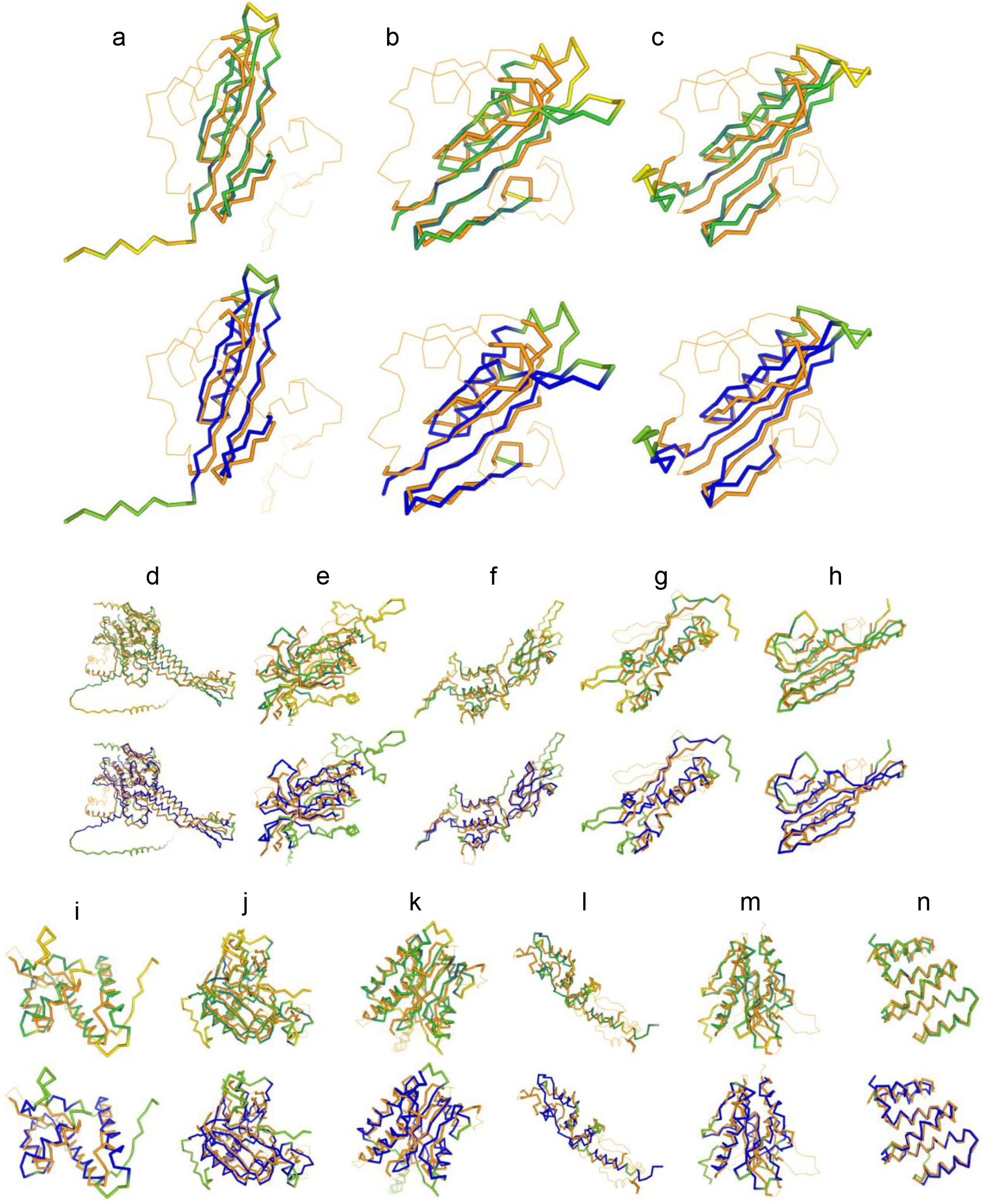
Structural homologs of CrV-01T cryptic proteins. In each panel, the top and bottom three-dimensional DALI superimpositions are colored by sequence and structure conservation, respectively. The C-α trace of each CrV-01T AlphaFold 3 model is colored using a yellow– green–blue gradient, yellow and green representing the lowest conservation values and blue the highest. The relative entropy was used as the conservation measure. The structural homologs are colored in orange. (**a**) Photosystem I reaction center subunit II (PsaD), open reading frames (ORF) 76 and 115. (**b**) PsaD, ORF 78. (**c**) PsaD, ORF 79 of CrV-like virus Cr-LKS4. (**d**) Portal protein, ORF 5. (**e**) Prohead core protein protease, ORF 8. (**f**) Head completion protein, ORF 22. (**g**) Multifunctional protein E4-ORF3, ORF 26. (**h**) Tail terminator protein, ORF 27. (**i**) Transcriptional regulator TnrA, ORF 47. (**j**) CRISPR associated protein, ORF 58. (**k**) Cyanobacteriochrome, ORF 65. (**l**) ArdA anti-restriction protein, ORFs 69 and 122. (**m**) Homing endonuclease, ORFs 80 and 111. (**n**) Peptidoglycan synthase activator, ORF 91.

CBCRs are widespread among cyanobacteria and play roles in regulating processes like phototaxis, cyclic nucleotide metabolism, and optimization of light harvesting^17^.

Although no photosystem structural homologs could be detected in Ma-LMM01, we were able to identify structural homology for twelve previously unannotated protein-coding genes (Fig. 2; Extended Data Table 2), all of which are transcribed during infection^18^. These included an IsrB nickase-like protein (ORF 171), which is a homolog of the Cas9 ancestor IscB^19^, and the putative central spike tip protein (ORF 75) of Ma-LMM01. We also found what appears to be the receptor-binding protein (RBP; ORF 87) of the virus, although only a region of the model shared structural homology with its DALI top hit, the RBP of *Lactococcus* phage 1358. As is sometimes the case for viral proteins^20^, both Ma-LMM01 and CrV-01T structural models were generally smaller compared to their cellular homologs. Notably, the models for which we were able to identify structural homology had a high and significant structural similarity (DALI Z-score > 4.2 in all cases) despite their very low amino acid sequence similarity (Figs. 1 and 2). The rest of the AF3 models were predicted with low confidence scores or did not share any clear structural homology with their DALI hits.

**Fig 2.**
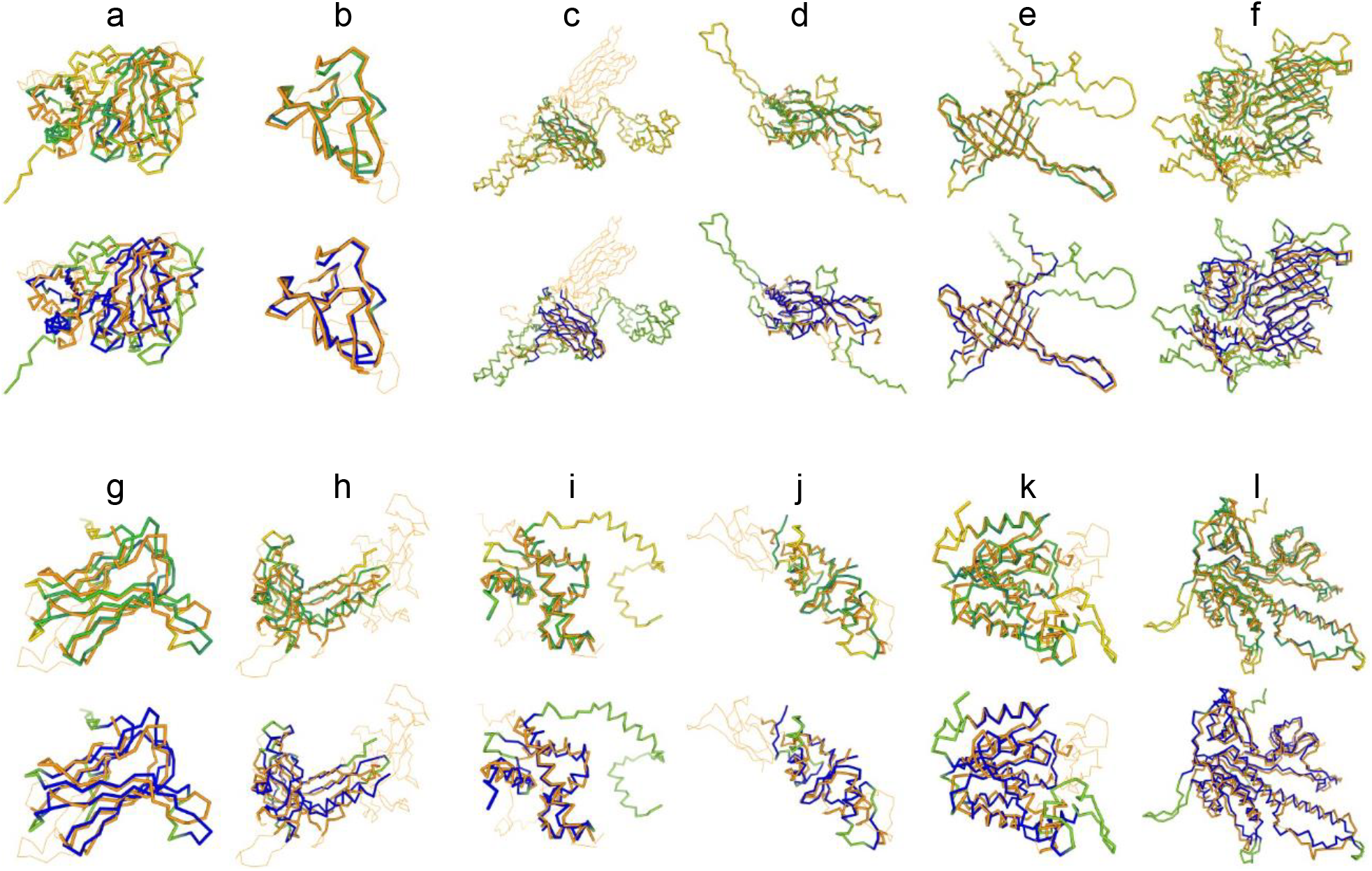
Structural homologs of Ma-LMM01 cryptic proteins. The structural models of the virus and their homologs are displayed and colored as in Fig. 1. (**a**) 7-carboxy-7-deazaguanine synthase, open reading frame (ORF) 72. (**b**) Central spike tip protein, ORF 75. (**c**) Receptor binding protein, ORF 87. (**d**) Tail terminator protein, ORF 90. (**e**) Tail tube protein, ORF 92. (**f**) Tail protein, ORF 93. (**g**) 5-hydroxy-isourate hydrolase, ORF 99. (**h**) Baseplate structural protein, ORF 116. (**i**) Q protein, ORF 124. (**j**) ArdA anti-restriction protein, ORF 158. (**k**) Adenylate kinase, ORF 162. (**l**) IsrB nickase, ORF 171.

So far, photosystem proteins have been considered to be more abundant in marine cyanophages than in freshwater ones^21^. Our results suggest that these proteins may be more common in freshwater cyanophages than previously thought. Photosystem protein-coding genes in freshwater cyanophages may have simply mutated to such an extent as to bear no detectable sequence similarity to their marine viral counterparts and their cellular homologs. The fact that we were only able to detect one sequence-based homolog of the CrV-01T putative PsaD proteins may hint towards highly divergent photosystem proteins among freshwater cyanophages as well. Indeed, lakes have long been proposed to work as island-like systems for evolutionary purposes^22^. However, it is still possible to identify these distant homologs using structural homology detection methods.

Overall, we have shown that previously isolated freshwater cyanophages contain multiple ORFs with structural homology to photosystem proteins, suggesting that they may be more widespread in these viruses than previously thought. Follow-up studies should investigate whether the transcriptional activity of the putative photosystem protein-coding genes from CrV-01T and Cr-LKS4 correlate with changes in the photosynthetic efficiency of the virocell; and apply AF3 proteomic modelling to a larger number of freshwater cyanophage genomes. The discoveries reported here emphasize the importance of applying structural homology detection methods when annotating viral genomes and highlight the potential of AlphaFold for exploring the dark matter of the aquatic virosphere.

## Methods

Proteome FASTA files containing all protein sequences encoded in the genomes of CrV-01T and Ma-LMM01, as well as ORF 79 of Cr-LKS4 were downloaded from the NCBI Protein database (GenBank accession numbers: MH636380, AB231700, and WHL30645, respectively). Structural modelling was conducted using AF3 through the AlphaFold server^7^. All protein-coding genes were modelled using automatically generated seeds and the default number of ten recycles. For each run (i.e., each three-dimensionally predicted amino acid sequence), AF3 outputs five structural models ranked by their predicted templated modelling (pTM) scores. In all cases, the top ranked models were used for downstream structural homology queries. The overall fold of AF3 models with a pTM score higher than 0.5 is usually considered similar to that of the true structure^23^. Thus, only the models with pTM scores higher than 0.5 were selected for further study. Structural homology queries were carried out using the PDB search tool of the DALI server^14^. Hits with a DALI Z-score > 2 (i.e., two standard deviations above expected) are usually considered significant^24,25^. Manual curation followed to further evaluate the relevance of the matches by visual inspection^26^. The single-chain structural alignments between the viral models and their structural homologs (Figs. 1 and 2) were generated and visualized using the 3D superimposition tool of DALI.

## Data availability

The data to reproduce the results of this study (i.e., the AF3 model seeds) are available in Extended Data Tables 1 and 2. Alternatively, the structural models can be provided upon reasonable request to I.M.P. The GenBank accession numbers for the previously sequenced genomes of CrV-01T and Ma-LMM01 are MH636380 and AB231700, respectively. The GenBank accession number for ORF 79 of Cr-LKS4 is WHL30645.

## Acknowledgements

This study would not have been possible without the continuous support of Dr. Esther Padilla Calderón and Dr. Medardo Meza Olea. This work was funded by a Consejo Nacional de Humanidades, Ciencias y Tecnologías (Conahcyt) Becas de Posgrado para Maestrías y Doctorados en Ciencias y Humanidades en el Extranjero scholarship awarded to I.M.P.; and Natural Sciences and Engineering Research Council of Canada (NSERC) Discovery Grants (2022-03350 and 2022-00329), and a Phycological Society of America Norma J. Lang Early Career Researcher Fellowship awarded to J.I.N.

## Author Contributions

I.M.P. conceived the idea, conducted the structural bioinformatics pipeline, analyzed the data and wrote the manuscript. A.C.D. provided additional insights on structural modelling. I.M.P. and

J.I.N. acquired funding. J.I.N. supervised the work. All authors revised and edited the manuscript.

---

**Extended Data Table 1.** Structural homologs of CrV-01T with no sequence-based functionally annotated homologs in the NCBI non-redundant protein sequences database.

| ORF | Sequence homolog<sup>a</sup> | BLAST *E*-value | AF3 pTM | Structural homolog | DALI Z-score | AF3 model seed |
| --- | --- | --- | --- | --- | --- | --- |
| 5 | 32 HPs | $< 1 \times 10^{-5}$ | 0.74 | Portal protein (Oshimavirus P7426) | 20.1 | 729767064 |
| 8 | 28 HPs | $< 3 \times 10^{-5}$ | 0.82 | Prohead core protein protease (T4) | 10.1 | 1619432911 |
| 22 | 8 HPs | $< 2 \times 10^{-7}$ | 0.68 | Head completion protein (T5) | 9.5 | 1459724406 |
| 26 | 73 HPs | $< 8 \times 10^{-4}$ | 0.88 | Multifunctional protein E4-ORF3 (Human adenovirus 5) | 4.7 | 68344560 |
| 27 | 13 HPs | $< 9 \times 10^{-7}$ | 0.88 | Tail terminator protein (*Rhodobacter capsulatus* GTA) | 10.5 | 1398212991 |
| 47 | 2 HPs | $< 9 \times 10^{-6}$ | 0.71 | Transcriptional regulator TnrA (*Bacillus subtilis*) | 5.8 | 873295830 |
| 58 | 9 HPs | $< 1 \times 10^{-6}$ | 0.87 | CRISPR associated protein (*Pseudomonas oleovorans*) | 15.8 | 1260485536 |
| 65 | 12 HPs | $< 4 \times 10^{-10}$ | 0.64 | Cyanobacteriochrome (*Nostoc punctiforme*) | 12.3 | 535717079 |
| 69 | ORF 72 of Cr-LKS4 | $2 \times 10^{-64}$ | 0.77 | ArdA anti-restriction protein (*Enterococcus faecalis*) | 4.9 | 1366043568 |
| 76 | ORF 79 of Cr-LKS4 | $5 \times 10^{-43}$ | 0.70 | PsaD (*Fischerella thermalis* PCC 7521) | 6.7 | 699591561 |
| 78 | N.a. | N.a. | 0.68 | PsaD (*Thermosynechococcus elongatus* BP-1) | 5.8 | 1900565996 |
| 80 | ORF 86 of Cr-LKS4 | $2 \times 10^{-97}$ | 0.83 | Homing endonuclease (*Synechocystis* sp. PCC 6803) | 9.3 | 234960824 |
| 91 | ORF 85 of Cr-LKS4 | $2 \times 10^{-42}$ | 0.84 | Peptidoglycan synthase activator (*P. aeruginosa*) | 16.4 | 1748855320 |
| 111 | ORF 86 of Cr-LKS4 | $2 \times 10^{-97}$ | 0.83 | Homing endonuclease (*Synechocystis* sp. PCC 6803) | 9.3 | 234960824 |
| 115 | ORF 79 of Cr-LKS4 | $5 \times 10^{-43}$ | 0.70 | PsaD (*F. thermalis* PCC 7521) | 6.7 | 699591561 |
| 122 | ORF 72 of Cr-LKS4 | $2 \times 10^{-64}$ | 0.77 | ArdA anti-restriction protein (*Enterococcus faecalis*) | 4.9 | 1366043568 |
| 79<sup>b</sup> | ORF 76 of CrV-01T | $5 \times 10^{-43}$ | 0.67 | PsaD (*F. thermalis* PCC 7521) | 6.7 | 514579827 |

Abbreviations: ORF, open reading frame; HPs, hypothetical proteins; AF3, AlphaFold 3; pTM, predicted template modelling score; GTA, gene transfer agent; PsaD, Photosystem I reaction center subunit II.
<sup>a</sup>As of December 1st, 2024. Only BLASTP hits with *E*-values $< 0.001$ were considered significant and counted.
<sup>b</sup>ORF from CrV-like virus CrLKS4.

**Extended Data Table 2.**
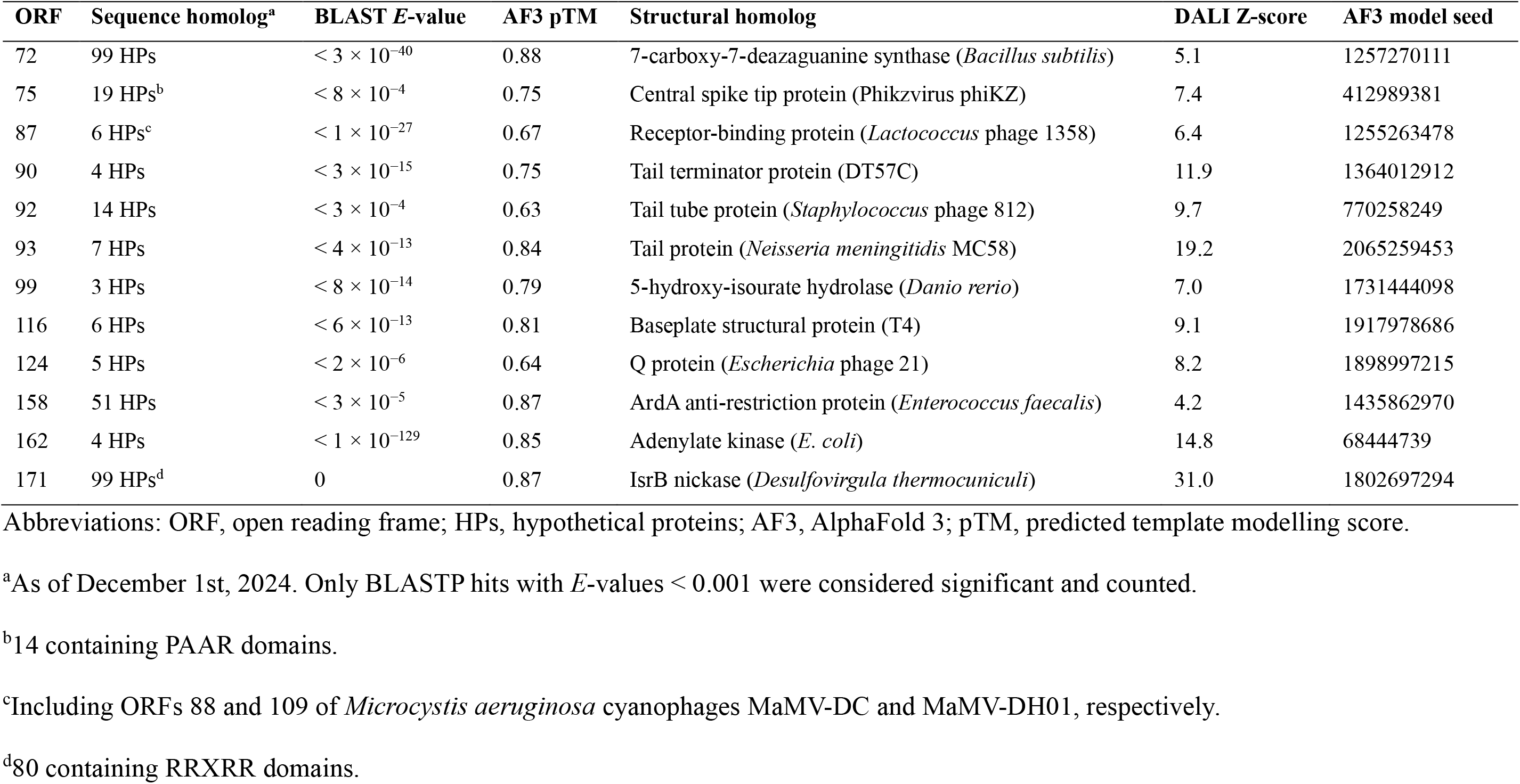
Structural homologs of Ma-LMM01 with no sequence-based functionally annotated homologs in the NCBI non-redundant protein sequences database.

